# Multimodal characterization of the murine neurovascular niche using a novel microvascular isolation protocol

**DOI:** 10.1101/2021.08.31.458349

**Authors:** Katrine Dahl Bjørnholm, Michael Vanlandewijck, Francesca Del Gaudio, Urban Lendahl, Per Nilsson, Helena Karlström, Christer Betsholtz

**Affiliations:** Department of Neurobiology, Care sciences, and Society, Division of Neurogeriatrics, Karolinska Institutet, Sweden; Department of Medicine, Huddinge, Karolinska Institutet, Sweden; Department of Immunology, Genetics, and Pathology, Rudbeck Laboratory, Uppsala University, Sweden; Department of Cell and Molecular Biology, Karolinska Institutet, Sweden

**Keywords:** Single cell RNA sequencing, microvascular isolation, neurovascular unit

## Abstract

The blood-brain barrier (BBB) is central to separate blood from the extracellular fluids of the brain. To understand disease-related changes in the BBB is pivotal and such changes can increasingly be studied by single-cell RNA sequencing (scRNAseq), which provides high-resolution insight into gene expression changes related to the pathophysiological response of the vasculature. However, analysis of the vascular cells in the brain is challenging due to the low abundance of these cells relative to neuronal and glial cells, and improved techniques for enrichment of the vascular component is therefore warranted. The present study describes a method whereby panning with CD31-coated magnetic beads allows isolation of brain vasculature without the need for transgenic reporter lines or FACS sorting. The protocol was tested in three modalities: isolation of cells for scRNAseq, western blot (WB) analysis, and primary cell culture. For scRNAseq, a total of 22,515 single-cell transcriptomes were generated from 12-months old mice and separated into 23 clusters corresponding to all known vascular and perivascular cell types. The most abundant cell type was endothelial cells (EC) (*Pecam1*- and *Cdh5*-positive), which dispersed into clusters of arterial, capillary, and venous EC according to previously established BBB arterio-venous zonation markers. Furthermore, we identified clusters of microglia (*Aif1-*positive), one cluster of fenestrated endothelial cells (*Plvap*-positive; *Cldn5*-negative), a cluster of pericytes (*Kcnj8*- and *Abcc9-*positive) and a cluster of vascular smooth muscle cells (VSMC) (*Acta2*- and *Tagln*-positive). WB analysis using established markers for the different cell types (CD31 (EC), SM22 (VSMC), PDGFRB (pericytes), GFAP (astrocytes), and IBA1 (microglia) confirmed their presence in the isolated vascular component and suggests that the protocol is suitable for future proteomic analysis. Finally, we adapted the isolation protocol to accommodate primary culture of brain vascular cells. In conclusion, we have successfully established a simple and fast method for isolating microvasculature from the murine brain independent of cell sorting and alleviating the need to use reporter mouse lines. The protocol is suitable for a multitude of testing modalities, including single-cell analyses, WB and primary cell culture.

## Introduction

Research into the organization of the brain vasculature and its response to injury and disease is a burgeoning research field. The mouse is increasingly used as an experimental model to study various aspects of brain vascular dysfunction, ranging from traumatic brain injury to neurodegenerative disease. Therefore, development of methodology allowing easy access to the mouse brain vasculature for experimental analysis is warranted.

The brain is an energy-demanding organ, which requires a complex vasculature to transport oxygen and nutrients to the brain. From cerebral arteries, blood continues via the pial arteries that reside in the subarachnoid spaces at the surface of the brain. From the pial arteries, penetrating arterioles branch off into the brain parenchyma and continue branching into capillaries deeper in the brain. Capillaries then join into venules and further into pial veins back into the subarachnoid spaces. The blood vessels are built from an inner tubing of endothelial cells, which are surrounded by mural cells (vascular smooth muscle cells (VSMC) and pericytes). In larger vessels, the endothelium is coated by VSMC while the capillaries are covered by pericytes (Lendahl et al., 2019). In addition to the “core” cell types of the vasculature – endothelial cells, pericytes and VSMC – there are also several other cell types associated with the brain vasculature. Astrocytes play an integral role in securing the blood-brain barrier (BBB), a structure which ensures that the blood is separated from the extracellular fluids of the brain. The astrocytic endfeet form a coverage that encapsulates the vasculature of the BBB. Furthermore, there are microglial cells, which are scavengers for extracellular debris and are activated in neuroinflammation. Finally, different types of perivascular fibroblasts are associated with the brain vasculature. There is an increasing realization that the brain vasculature undergoes considerable changes in response to injury or brain disease, for example following traumatic brain injury, stroke and cerebral small vessel disease. There is also mounting evidence for the importance of a dysfunctional brain vasculature in neurodegenerative diseases, including Alzheimer’s disease.

To understand how the brain vasculature functions under normal homeostatic conditions and becomes dysfunctional upon brain injury and disease, it is important gain insights into the molecular changes in the various cell types. Single cell RNA-sequencing (scRNA-seq) has emerged as a revolutionizing technology to decode gene expression changes at the single cell level and is increasingly applied also to the brain vasculature(Vanlandewijck, He et al. 2018, Zeisel, Hochgerner et al. 2018). However, the study of vascular cells in the brain is challenging due to the low abundance of these cells relative to neuronal and glial cells (Saunders, Macosko et al. 2018). Previous investigations have successfully used reporter mice strains for the cells of the neurovascular unit and isolated the cells using FACS (Vanlandewijck, He et al. 2018). The generation and use of mouse reporter strains is however a long and cumbersome process especially if investigations are to be carried out in several different *in vivo* animal models and if these models need to be crossed into various disease models. Moreover, the FACS sorting procedure to capture the cell type of interest from a transgenic reporter strain tends to lower cell integrity and viability. To overcome these current experimental hurdles, we have developed a magnetic bead protocol that enables isolation of microvascular segments and applied this to single cell sequencing investigations as well as protein chemistry and cell culture investigations. This enables analysis of the vasculature on single cell level in any mouse model, alleviating the need for the use of mouse reporter cell lines and FACS sorting prior to analysis.

## Materials and methods

### Mice

C57Bl/6J mice were purchased from Jackson Laboratory (Bar Harbor, ME, USA). Three male 12 months old mice were used for the generation of both single cell and western blot data. Another three mice were used to complement the western blot data (for a total of 6 samples, see figure 3) Four mice of mixed sex were used for in vitro cell culture. The reporter mice used in the *in vitro* studies (Pdgfrb-eGFP and Ng2-DsRed report mice) are described earlier (Vanlandewijck, He et al. 2018). All studies were approved by Stockholm ethical committee for animal experiments (6-14, 1433-2018, 12370– 2019), Stockholm North Ethical Committee on Animal Research (Permit number N16/12), Linköping ethical board for animal experiments (ID407) and by the Uppsala Ethical Committee on Animal Research (Permit number: C224/12 and C225/12). All experimentation was carried out by licensed personnel. All mice were housed in cages of 3-5 per cage under 12 h light dark cycle at 22°C and 50% humidity with *ad libitum* access to food and water. Animal caretakers inspected animals daily.

### Microvascular isolation

Mice were euthanized by cervical dislocation and the head was immediately removed and placed in ice cold primary washing medium (Dulbecco’s Modified Eagle Medium (DMEM) with 1% Penicillin/Streptomycin (P/S), both from Gibco. Once in the lab, the brain was carefully dissected from the skull and the brain stem and olfactory bulbs were removed. For scRNA-seq, the cerebellum was also removed and only one hemisphere was used for further analysis. The brain was then placed in a petri dish and minced by cutting with a pair of scissors parallel to the petri dish plane to avoid cutting into the plastic surface (<1mm^3^ pieces). Once homogenous, the tissue was dissolved in 3 ml freshly prepared primary dissociation buffer (collagenase 1 mg/ml, Sigma C6885, in DMEM w/o phenol red, Gibco) and passed through a cut p1000 pipet tip ten times or until smooth passing. The solution was then transferred to a 5 ml Eppendorf tube and placed in a rotating incubator (speed 20 rev/min) at 37°C for 10 minutes. Samples were then passed through an uncut p1000 pipet tip 10 times or until smooth passing. Samples were then placed back in the rotating incubator for 10 minutes at 37°C. The samples were filtered through a 70 µm mesh strainer (Falcon) into a 50 µl tube (Falcon) to remove myelin. The filter was washed with primary washing medium and samples were spun at 300 g for 5 minutes at RT. The supernatant was removed, and the pellet was resuspended in 4.5 ml secondary washing medium (DMEM with 1% PS (Gibco), 0.5 mg/mL heparin (Sigma)) and divided into three 2 ml Eppendorf tubes. 15 µl Rat anti-mouse CD31 antibody (BD, 10 µl/brain) coupled Dynabeads^®^ Sheep Anti-Rat IgG (Invitrogen, 50 µl/ brain, prepared according to manufacturer’s instructions) were added to each subsample and placed in rotating incubator for 30 minutes at room temperature. After antibody incubation the tubes were placed on a DynaMag 2 magnet (Invitrogen), and the supernatant was removed. Beads were washed twice by resuspension and magnetic capture in secondary washing medium, before the three samples were pooled in one 2 ml Eppendorf tube. The resuspended beads coupled with microvasculature were visualized using an inverted microscope at 20x magnification. Dissociation of the microvascular fragments from the beads was done in different ways, depending on the downstream application, as detailed below.

### Single cell capture and cDNA generation

For single cell sequencing experiments and western blot, the supernatant was replaced with 1.5 ml freshly made secondary dissociation buffer (10xTrypLE with 1 mg/ml collagenase IV, both from Gibco). Samples were placed in rotating incubator for 5 minutes at 37°C then passed through a 20G needle with a 1 ml syringe and placed back in rotating incubator (37°C) for 5 minutes. Beads were captured on the magnet and the supernatant now containing the dissociated cells was transferred to a 15 ml Falcon tube containing 10 ml ice cold FACS buffer (DMEM w/o phenol red + 1% P/S, 2% FBS, 1 mM EDTA pH 8, all from Gibco) and centrifuged for 5 minutes at 300 g and 4°C. The pellet was resuspended in 100 µl FACS buffer and cells were counted using the Luna FL dual fluorescence cell counter (Logos biosystems).

After counting, single cells were captured using Next Gem Single Cell 3’ Reagent kit v3.1. (Dual index) from 10x Genomics. Samples were loaded to capture 5,000 cells as per manufacturers instruction. Full libraries were sequenced using Illumina platform on a Next seq2000 P3 flow cell, aiming for an average read depth of 40,000 reads/cell.

### Single cell Data analysis

Flow cell data were demultiplexed and aligned using the cell ranger pipeline (10X genomics) and the Seurat package version 4.0.1 (Hao, Hao et al. 2021) for R version 4.0.5. Clustering analysis was done by PCA reduction in 17 dimensions at 0.8 resolution.

### Western blot

After the isolation, 15 μl of cell suspension (same as used for single cell RNA sequencing) was mixed 1:1 with sample buffer (62.5 mM Tris-HCl, pH 6.8, 25% glycerol, 2% SDS, 5% β-mercaptoethanol, 0.01% bromophenolblue), heated for 10 min at 95°C and loaded on Mini Protean TGX precast gel 4-12% (Bio-Rad). For the blot, Trans-Blot Turbo nitrocellulose membrane transfer pack (Bio-Rad) was used. The membranes were blocked with StartingBlock™ (PBS) Blocking Buffer (Invitrogen) for 30 min at room temperature with gentle agitation. Primary antibodies (anti-CD31 #77699, anti-Pdgfrβ #3695 from Cell Signaling; anti-GFAP #63893 and anti-β-actin #A5441 from Sigma Aldrich; anti-Sm22 #14106 from Abcam and anti-Iba1 #01919741 from Wako) were diluted 1:1000 and incubated in 1:1 Blocking buffer and PBS-0,1% Tween20 overnight at 4°C with gentle agitation. Then, the membranes were washed in PBS-0,1% Tween20 three times for 15 min at room temperature and anti-rabbit or anti-mouse secondary antibodies (1:3000, Cell signaling) were applied for 30 min at room temperature with gentle agitation. After three additional washes, the antibody signal was captured with Super Signal West PICO Plus Chemiluminescence Substrate (Thermo Fisher) using Gel Doc (Bio-Rad). Each sample represents a wild-type mouse. Six samples were used for this experiment.

### Cell culture

For cell culture of microvessels, the bead-coupled microvasculature was resuspended in 1.5 ml of pure 10xTrypLE buffer for 5 minutes at 37°C without agitation. Then, the samples were inverted 5 times by hand, after which they were incubated for another 5 minutes at 37°C. This step ensures dissociation of microvascular fragments from the beads. After this step, the samples were placed on the magnet again to remove the dissociated beads. The supernatant was transferred to a 15 ml Falcon tube and an excess of ice-cold FACS buffer (12 ml) was added to the samples, after which they were spun down for 5 minutes at 300 g and 4°C. The supernatant was removed, and the vascular fragments were seeded in wells of a 24-well plate, pre-coated with collagen IV (Sigma-Aldrich: C6745) at 10 µg/cm^2^, as per the manufacturer’s recommendation. Microvascular fragments from one brain were seeded in 6 wells of a Corning^®^ 24-well plate or a single well of a Corning^®^ 6-well plate. For live and confocal imaging, glass-bottom plates (Mattek Corp.) were used. Vascular fragments were cultured in Endothelial Cell Growth Medium 2 (PromoCell: C22011) with full supplement and 1% Penicillin/Streptomycin. Cell medium was changed every 24 hours. Cell cultures were not split upon high confluency, as this was detrimental to the culture (data not shown).

### Live cell imaging

Vascular fragments were prepared and seeded on glass bottom 24-well plates as described above. Cells were imaged in an Operetta CLS High-Content Analysis System (PerkinElmer) with one image taken every 15 minutes for 72 hours. Every 24 hours, the cell medium was changed. To create movies from single images, a custom Java Script was used to organize the images in sequential order. Afterwards, movies were generated at 25 images per second using the Fiji distribution of Imagej2 (https://imagej.net/software/fiji/). Live cells were imaged using a Leica DMIL microscope equipped with a Lumencore LED light source and a Leica DFC 450 C camera.

### Confocal imaging

Vascular cell cultures were seeded on glass bottom 6-well plates as described above. The cells were fixed and stained with their respective antibodies as described before (Vanlandewijck, He et al. 2018). Image acquisition was done on a Leica SP8 confocal microscope system equipped with a White Laser. Image processing was done using Adobe Photoshop CS6 or Fiji.

## Results

### Recovery and viability of vascular cells

After first digestion of a single hemisphere with collagenase II and subsequent filtering through a 70µm filter, the vascular stubs were captured with CD31-coupled Dynabeads (Fig 1A). Visualization in an inverted microscope showed an abundant number of beads, bundled together in rod-shaped structures identified as vascular stubs (Figure 1B, black arrows). After a second digestion step, designed to release the beads and further dissociate the microvascular segments into single cell suspension, the cells were counted. Over a range of 48 samples, a mean viability of 92.1% live cells (SEM +/- 0.71, Figure 1C) and a mean recovery was 1474 cells/µL (SEM +/- 99.7, Figure 1D) was obtained. All samples were resuspended in 100µL of FACS buffer, resulting in a total cell recovery range of single cells between 54,900 and 347,000 cells from a single hemisphere (Fig 1F).

**Figure 1:**
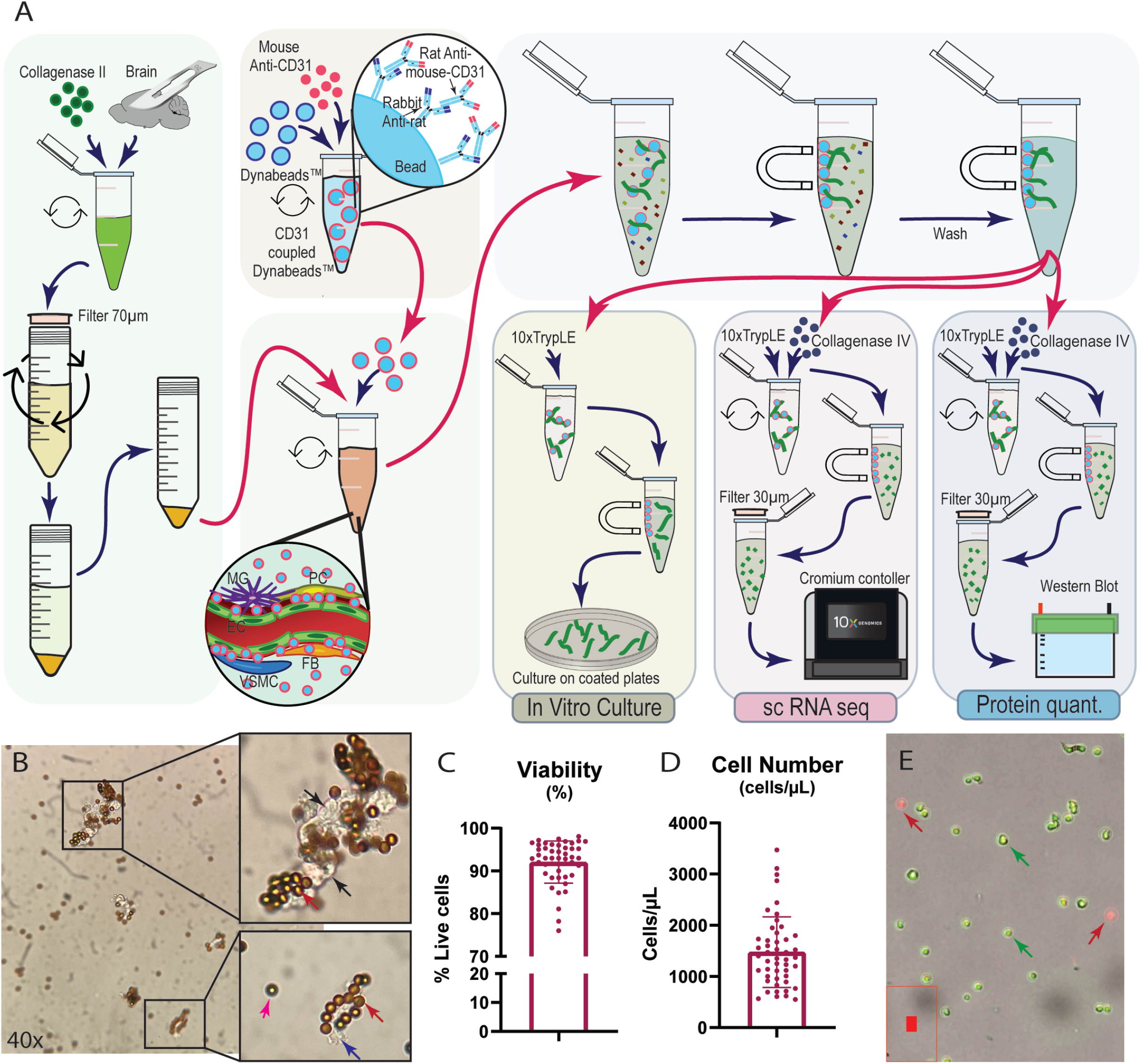
Methods. **A**. Simplified illustration of the steps in the method for microvascular isolation. Blue arrows indicate changes within the same step. Red arrows signify the beginning of a new step. **B**. Light microscopy image of microvascular fragments coupled to magnetic beads (40x magnification). Top insert shows a large fragment (black arrows) and coupled magnetic beads (red arrow). Bottom insert shows a smaller vascular fragment (blue arrow) with coupled beads (red arrow) and a free-floating bead (pink arrow). **C**. Viability score (n=42), **D**. Cell number per µL (n=42). E: Image showing single cell suspension stained for viability (dual stain Acridine Orange/Propidium Iodide) Live single cells are green (green arrows) and dead cells in red (red arrows). Picture from Lunar FL cell counter software, 2x magnification. scRNA seq: single cell RNA sequencing, EC: endothelial cells, MG: microglia, PC: pericytes, VSMC: vascular smooth muscle cells, FB: fibroblasts, Protein quant.: protein quantification.

### Single cell sequencing of microvasculature

Next, vascular isolation was performed on three C57Bl6 mice of 12 months old and the cells were analyzed as single cells using the 10x Genomics gene expression chemistry (v3.1). A Uniform Manifold Approximation and Projection (UMAP) representation of all the cells is shown in Figure 2A. From the three samples, 23,582 cells were isolated and after filtering (nFeature_RNA > 200 & nFeature_RNA < 5000 & percent.mt < 10, Figure 2B) 22,515 cells were included in the analysis. Data clustered into 23 clusters (Figure 2A), which were annotated using the top ten most expressed genes (Figure S1).

**Figure 2:**
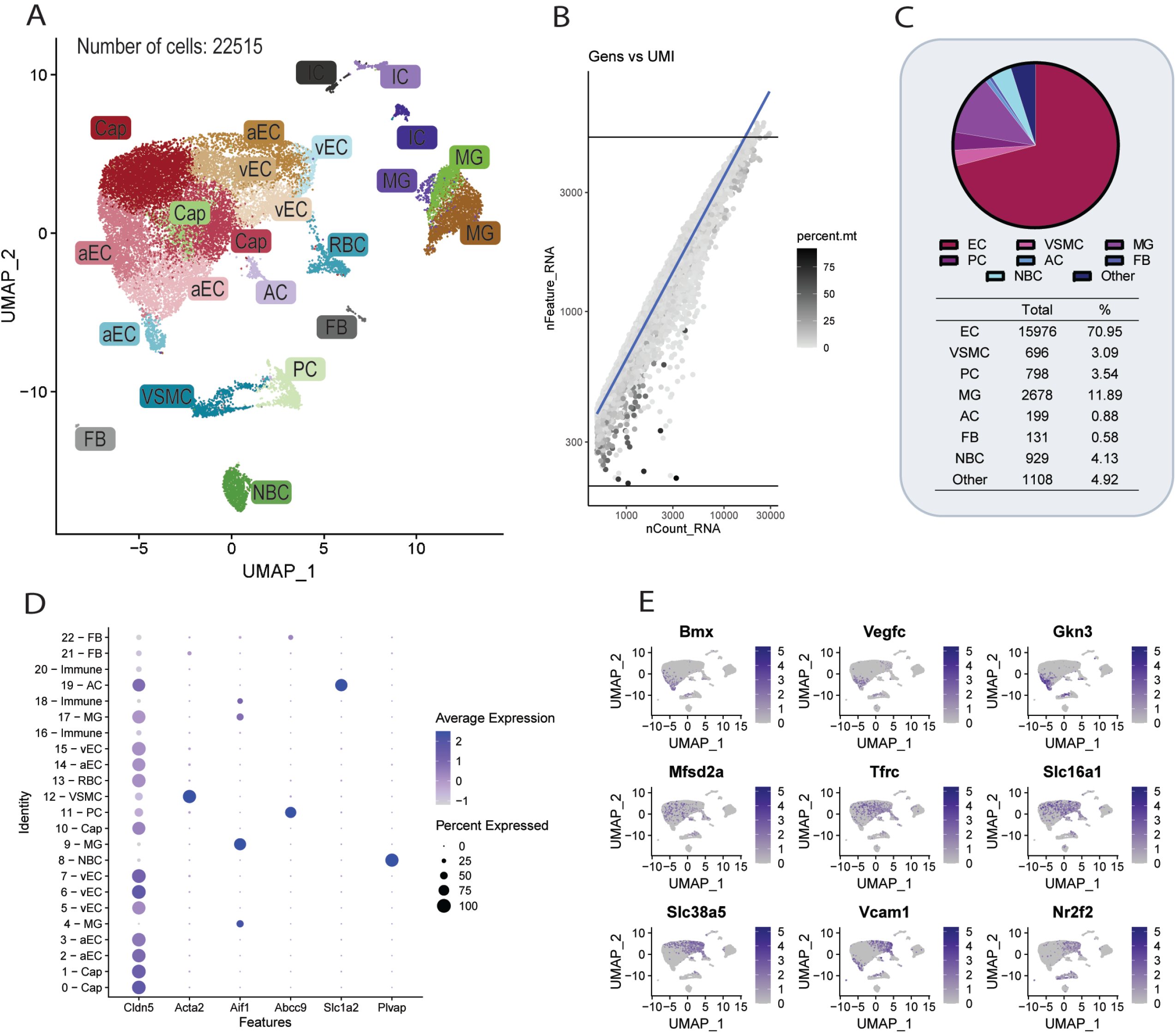
Single cell sequencing. **A**. UMAP showing how the 22515 cells from three 24 months old WT mice distribute intro 23 clusters. **B**. Plot showing the quality of cell after sequencing by number of unique molecular identifiers (UMI, nCount_RNA) as a function of number of genes detected (nFeature_RNA). Color signifies percent mitochondrial RNA (percent.mt). **C**. Pie chart showing the distribution of the different cell types in the isolated cells. Below the chart, a table of cell count for each cluster and respective percentages. **D**. Dot plot showing cluster specific markers used for annotation of clusters. **E**. Grid of feature plots showing the endothelial cell zonation by markers of arterial endothelial cells (top row), capillary endothelial cells (center row), and venous endothelial cells (bottom row). AC: astrocytes, aEC: arterial endothelial cells, Cap: capillary endothelial cells, EC: endothelial cells, FB: fibroblasts, IC: immune cell, MG: microglia, NBC: non-barrier cells, PC: pericyte, RBC: red blood cells, vEC: venous endothelial cells, VSMC: vascular smooth muscle cell,

The most abundant cell type was endothelial cells (70.95%, Figure 2C) characterized by expression endothelial markers including *Cldn5* and *Cdh5* (Figure 2D). The endothelial cells divided into 13 subclusters encompassing venous, capillary, and arterial endothelial cells, confirming the previously established arteriovenous zonation of the blood brain barrier (Vanlandewijck, He et al. 2018) (Figure 2E). The second most abundant cell type was microglia (MG, 11.89%), based on the expression of specific MG markers including but not limited to *Aif1*. The MG separated into three subclusters. Moreover, *Acta2* was highly expressed in 100% of cells in cluster 12, which was identified as vascular smooth muscle cells (VSMC) (3.09%, Figure 2C). Cluster 11 expressed the Pericyte (PC)-specific marker *Abcc9* and this cluster amounted to 3.54% of the total amount of cells. A cluster of *Plvap*-expressing cells were identified as non-barrier endothelial cells (4.12%), possibly arising from choroid plexus. A small cluster of *Slc1a2*-expressing cells was identified as astrocytes (AC, 0.88%), and two small but distinct clusters were identified as fibroblasts (FB, 0.58%). Furthermore, four clusters, amounting to 4.92% of the total dataset, were identified as either red or white blood cells.

### Verification of microvascular cell clusters on the protein level

To confirm the presence of the identified clusters, Western Blot analysis was performed on isolated microvessels from six different mice (Figure 3). For endothelial cells, CD31 was used as marker, while Sm22 was used as marker for VSMC. To confirm the presence of pericytes, Pdgfrb was used. Astrocytes were visualized by GFAP as marker and lastly, Iba1 was used as a marker for microglial cells. B-actin was used as a loading control, which varies between samples due to the difference in isolation efficiency between experiments. However, the presence of the cell types expected from the scRNAseq experiments was confirmed.

### Cell culture of microvascular fragments

As an extension of the microvascular isolation protocol, a cell culture protocol was developed with the aim to establish a tool for *in vitro* study of mouse brain vasculature. In this protocol, the cells are not dissociated into single cells when the coupling of the fragments to the beads is disrupted. Instead, a milder dissociation is used to preserve the microvessel architecture when seeding the cells. After an investigation of several substrates, collagen IV was selected as the culture substrate of choice, as it provided the best cell survival. To be able to trace the cells, microvasculature from reporter mice expressing Ng2-DsRed were used.

To verify the results from the scRNAseq experiment, markers for the most common (>1% representation) cell types in the microvascular fragments were used for immunohistochemistry (Figure 3A). Indeed, the cell culture comprised of mostly endothelial cells (as visualized with anti CD31 antibodies). In addition, pericytes were identified that were positive for both CD13 and the Ng2-DsRed reporter of the mouse used for extraction of the cell culture. Surprisingly, staining for smooth muscle cell markers provided weak to negative staining. For technical reasons, staining for microglial cells was not performed. In order to confirm the presence of pericytes in the cell culture, cells were isolated from a dual reporter Pdgfrb-eGFP x Ng2-dsRed mouse. Indeed, after 72 hours of culture, cells with a clear pericyte morphology could be observed (Figure 3B).

**Figure 3:**
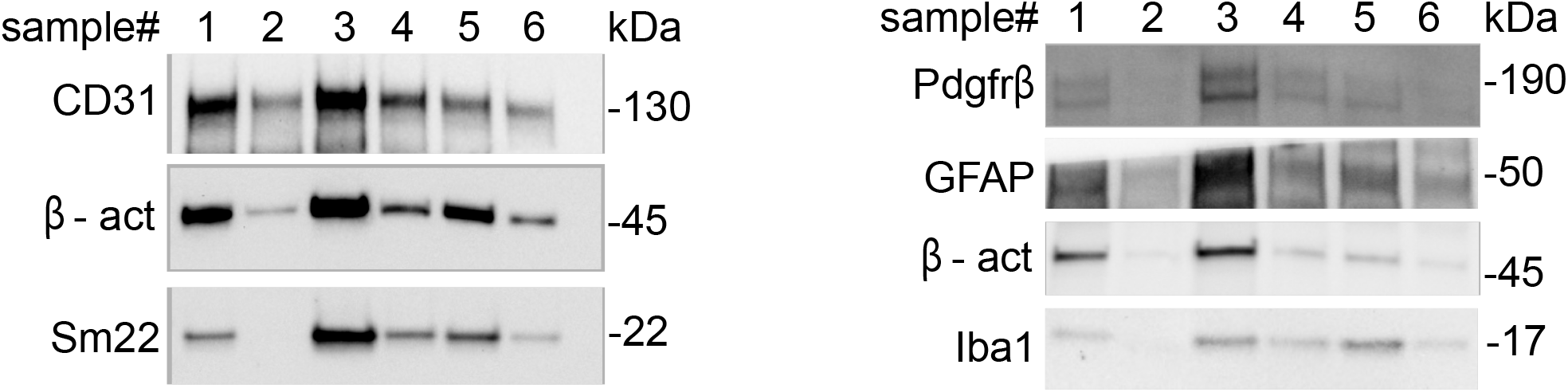
Western blot for markers of the single cell clusters. CD31 is used as a marker for endothelial cells, while Sm22 is specific for smooth muscle cells. Pdgfrb is expressed in all mural cells. GFAP and Iba1 are makers for astrocytes and microglial cells, respectively. β-actin is used as a loading control. The western blot is a representative example of three independent experiments.

**Figure 4:**
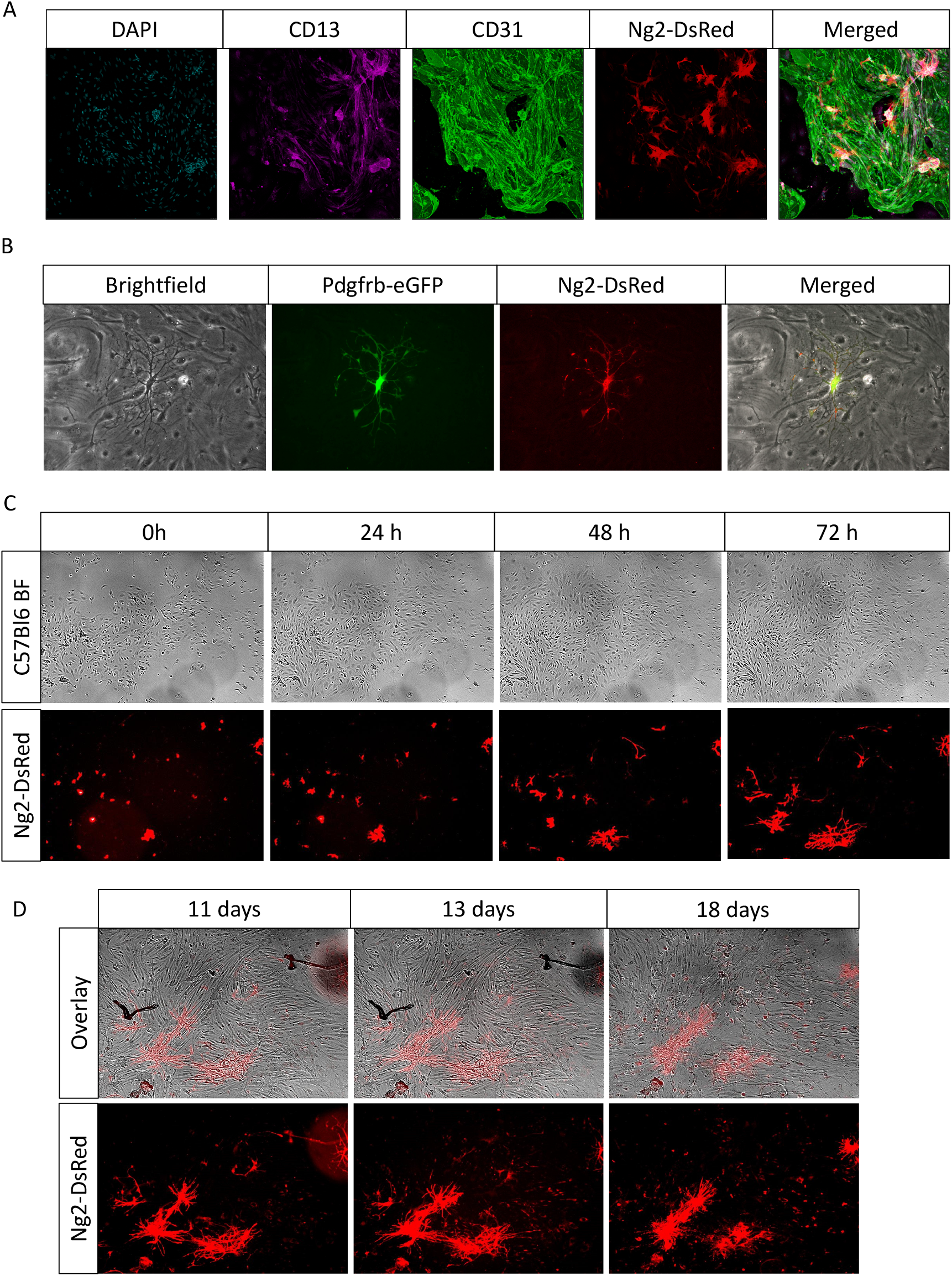
Cell culture of microvascular fragments. **A**. Confocal imaging of a microvascular culture 48 hours after seeding, isolated from an Ng2-DsRed mouse. DAPI is used to visualize the nuclei. CD13 is a marker for mural cells. CD31 visualizes the endothelial cells and Ng2-DsRed is a reporter for mural cells. **B**. Imaging of microvessel cell cultures 72 hours after seeding. The cells were isolated from a Pdgfrb-eGFP x Ng2-Dsred reporter mouse. Note the clear pericyte morphology on top of the endothelial cells. **C**. Still images from a live cell growth recording. Note that the top panel displays cells isolated from a WT C57bl6 mouse, while the bottom panel contains cells isolated from an Ng2-dsRed reporter mouse. The movie from which these images are extracted is found in Supplementary movies 1 and 2. Note that the time scale indicates time from imaging start, which is done 24 hours after seeding. **D**. Imaging of the same cell culture as in the bottom panel of (C), 11, 13 and 18 days after seeding. Note the excessive cell death and loss of pericyte processes 18 days after seeding.

Finally, to investigate the spatiotemporal dynamics and longevity of cells in the cell culture system, live imaging of isolated microvessels was performed (Figure 3C/D and supplementary movies 1 and 2). The imaging was started 24 hours after cell seeding to allow the cells to recover from the isolation procedure and attach firmly. At early stages, clusters containing pericytes centrally could be observed. These are likely originating from microvascular fragments that contained an attached pericyte. Over time, the endothelial cells patches expand, and tip cell migration can be observed (Supplementary movie 1). The pericytes, on the other hand, did not migrate, yet attempted to cover as many endothelial cells as possible by extending radial protrusions. It was also noted that endothelial cells covered with pericytes did not migrate or divide, in line with *in vivo* observations that pericytes stabilize vessels and preserve their maturity (Mae, He et al. 2021). Lastly, after extending the cell system to longer time periods, we observed that around 18 days after isolation, excessive cell death as well as disconnection of pericyte arms took place.

## Discussion

Primary isolation of brain vasculature is paramount for several types of analyses. However, isolation protocols are often lengthy (compromising cellular identity), require a lot of input material, and/or require the use of transgenic mouse reporter strains. In this report, we present a fast and efficient isolation protocol for microvascular fragments which is applicable to many downstream analyses. Our protocol is highly reproducible and isolates all components from the neurovascular unit, albeit not in the proportions that are found *in vivo*. For example, roughly 70% of our dataset is comprised of endothelial cells, while pericytes only encompass 3,5% of the dataset. This is clearly a deviation from the expected one pericyte cell body for every six endothelial cells ratio (Mae, He et al. 2021), and is most likely due to the use of CD31, an endothelial marker, as the tool for microvessel capture. Conversely, the proportionally low number of astrocytes is likely explained by the relative long distance of the astrocyte cell bodies from the vessels (Dubois, Campanati et al. 2014). Fibroblast-like cells, which are present in low numbers in the present dataset, are known to be associated with the arteries and arterioles (Vanlandewijck, He et al. 2018), and with this protocol it is likely that we capture arterioles and capillaries mainly, explaining the relative low number of fibroblast-like cells. The same explanation is probably relevant for the low number of VSMC. Minimizing enzymatic digestion of the whole brain tissue might result in more mural cells adhering to the endothelial cells; however, this introduces the risk of filtering away larger fragments, compromising cell recovery all together. Furthermore, it increases the risk of insufficient suspension into single cells, which is pivotal for generating high-quality single cell data. The data from the scRNAseq experiments were corroborated by WB analysis, as the expected cell types could be identified using cell type-specific antibodies. The WB data also indicate that proteins can be obtained without obvious degradation, which opens vistas for conducting proteomics-based studies, using this protocol. Vascular isolation from a single hemisphere consistently yielded enough material for scRNA-seq and WB analysis, opening up for the possibility to correlate gene expression to protein levels within the same samples.

The protocol also allows for establishing in vitro cultures of the brain vasculature, and the principal vascular cell types could be identified by immunostaining, although VSMC were under-represented, which likely is a consequence of that the culture conditions we employed were aimed for maximal recovery and preservation of endothelial cells. We observed pericyte attachment to endothelial cells, but it yet remains to be fully explored how well the three-dimensional vessel structure is retained in the in vitro cultures. Notably, we observed that pericytes displayed a different morphology as compared to *in vivo*. While *in vivo*, pericytes display a longitudinal extension along the vessel in both directions, in our in vitro cultures, the pericytes present numerous protrusions. This is likely explained by the absence of a three-dimensional vessel structure in the culture system, provoking the pericyte to cover as many endothelial cells as possible. This observation also goes in line with the observation that pericytes elongate substantially in pericyte-deficient mouse models, such as the Pdgfb^ret/ret^ model (Armulik, Genove et al. 2010).

In conclusion, we provide here a robust, simple and rapid protocol that maximizes cell viability and gain for isolation of the principle cell vascular and vasculature-associated cell types of the mural brain. Since our method does not require the use of reporter lines to be bred into the disease model of study, it simplifies breeding strategies, reduces the usage of mice and leads to faster high-quality results. Furthermore, while this protocol has been optimized for the isolation of brain microvasculature, it can be easily adjusted for the isolation of vasculature of other tissues. Indeed, CD31 is expressed in endothelial cells of many organs, and as such, with minor modifications, can be used for isolation beyond the murine brain.

## Acknowledgement

We thank Dr. Colin Niaudet and Dr. Konstantin Gängel for initial assistance and advice in setting up the isolation protocol. We thank Jürgen Cammaerts for coding of the software tool for movie generation. We also would like to thank the core facility at Novum, BEA, Bioinformatics and Expression Analysis, which is supported by the board of research at Karolinska Institutet and the research committee at the Karolinska hospital. The financial support from Hjärnfonden (PN, UL and CB) is gratefully acknowledged.

## Figure legends

**Supplementary Figure 1:**
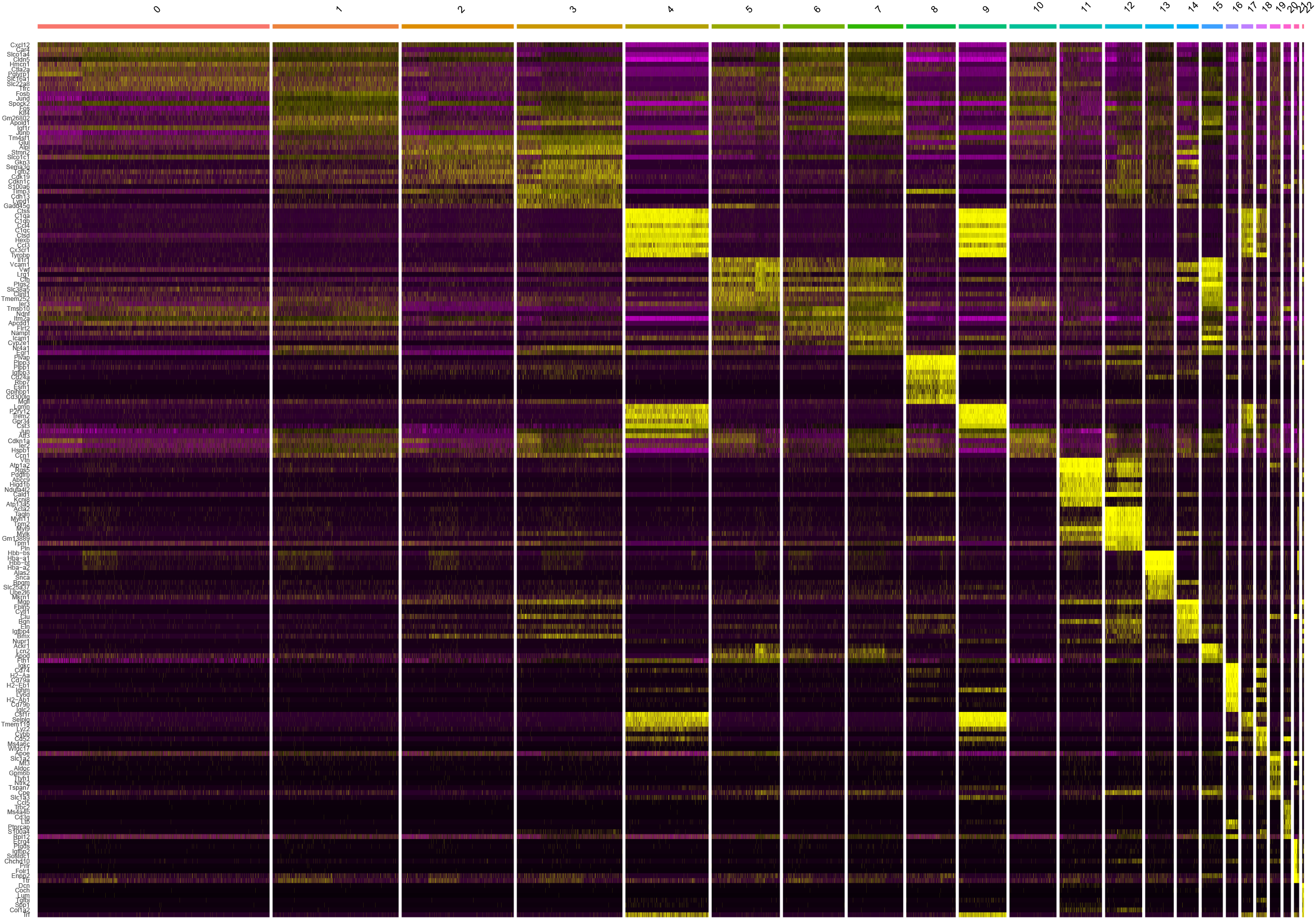
Heat map of top expressed gene per cluster. For every cluster, the ten most differentially expressed genes are displayed. Every column is a cell. Yellow indicates high expression, while black indicates low/absent expression.

**Supplementary movie 1: Live cell imaging of primary isolated microvessels of a C57Bl6 mouse**

The movie starts 24 hours after seeding and lasts for 72 hours of growth.

**Supplementary movie 2: Live cell imaging of primary isolated microvessels of a Ng2-dsRed reporter mouse**.

The movie starts 24 hours after seeding and lasts for 72 hours of growth.

